# Osmotic stress in roots drives lipoxygenase-dependent plastid remodeling through singlet oxygen production

**DOI:** 10.1101/2024.04.22.590524

**Authors:** Dekel Cohen-Hoch, Tomer Chen, Lior Sharabi, Nili Dezorella, Maxim Itkin, Gil Feiguelman, Sergey Malitsky, Robert Fluhr

**Affiliations:** Department of Plant and Environmental Sciences, Weizmann Institute, Rehovot 76100, Israel; Department of Chemical Research Support, Weizmann Institute of Science, Rehovot 76100, Israel; Department of Life Sciences Core Facilities, Weizmann Institute of Science, Rehovot 76100, Israel

**Keywords:** Defense response signaling mechanisms, cell fate, organelles, membrane mobility, *Arabidopsis thaliana*, root physiology

## Abstract

Osmotic stress, caused by the lack of water or by high salinity, is a common environmental problem in roots. Osmotic stress can be reproducibly simulated with the application of solutions of the high-molecular-weight and impermeable polyethylene glycol. Different reactive oxygen species such as singlet oxygen, superoxide and hydrogen peroxide accompany this stress. Among them, singlet oxygen, produced as a byproduct of lipoxygenase activity, was shown to be associated with limiting root growth. To better understand the source and effect of singlet oxygen, its production was followed at the cellular level. Osmotic stress initiated profound changes in plastid morphology and vacuole structure. By confocal and electron microscopy the plastids were shown to be a source of singlet oxygen accompanied by the appearance of multiple small extraplastidic bodies that were also an intense source of singlet oxygen. A marker protein, CRUMPLED LEAF, indicated that these small bodies originated from the plastid outer membrane. Remarkably a type 9 lipoxygenase, LOX5, was shown to change its distribution from uniformly cytoplasmic to a more clumped distribution together with plastids and the small bodies. In addition, oxylipin products of type 9 lipoxygenase increased while products of type 13 lipoxygenases decreased. Inhibition of lipoxygenase by SHAM inhibitor or in down-regulated lipoxygenase lines prevented cells from initiating the cellular responses leading to cell death. In contrast, singlet oxygen scavenging halted terminal cell death. These findings underscore the reversible nature of osmotic stress-induced changes, emphasizing the pivotal roles of lipoxygenases and singlet oxygen in root stress physiology.

## Introduction

Osmotic stress is one manifestation of drought stress. Proper osmotic adjustment in the plant is critical for maintaining crop yield under drought stress (Blum, 2017). Cells in the root function as an osmotic sensory organ and use various subcellular components to help maintain homeostasis. For example, vacuoles through the function of the tonoplast store a variety of osmolytes, to balance internal and external osmotic pressures. The accumulation of these solutes in the vacuole, mediated by specific transporters in the tonoplast increases the vacuole’s osmotic pressure, drawing water into the cell and maintaining turgidity essential for plant growth and stability (Maurel et al., 2015).

Plastids adjust their metabolic pathways to produce osmoprotectants, which are molecules like proline and sugar alcohols that help stabilize proteins and cellular structures, and maintain cell turgor (Hare et al., 1998). In addition, plastids play a role in the synthesis of abscisic acid (ABA), a critical hormone that regulates stress adaptation (Cutler et al., 2010). Plastids also play a role in generating reactive oxygen species (ROS) reactive oxygen species (ROS) accumulate within minutes after cell stimulation (Fichman and Mittler 2021). When the levels of ROS rise within cells, they can cause cellular destruction through lipid peroxidation, protein, RNA and DNA oxidation and are detoxified by antioxidant mechanisms (Dietz et al., 2016; Miller et al., 2010). However, ROS production coupled to regulated redox change in receptor proteins can be part of cellular and tissue signal transduction pathways.

Chiefly prominent in ROS synthesis during osmotic stress are NADPH oxidases of the RESPIRATORY BURST OXIDASE HOMOLOG (RBOH) family that produce superoxides that dismutate to hydrogen peroxide (Ben Rejeb et al., 2015). The activation of RBOH in stress depends on phosphorylation through e.g. CALCIUM-DEPENDENT PROTEIN KINASE 5 (Dubiella et al., 2013). Accumulation of ROS was shown to be associated with the clustering of PLASMA MEMBRANE INTRINSIC PROTEIN2;1 aquaporin (PIP2;1) that may impact directly on osmoregulation (Martinière et al., 2019).

Root tissue subjected to drought and osmotic stress has been shown to produce singlet oxygen in addition to other ROS (Chen and Fluhr, 2018; Mor et al., 2014). However, only singlet oxygen was correlated with a decrease in root growth, whereas other ROS do not exhibit this relationship (Chen and Fluhr, 2018). Indeed, a substantial transcriptomic correlation was noted between multiple abiotic stress scenario and singlet oxygen (Mor et al., 2014). These abiotic stresses include dryness, salinity, injury, and in certain conditions exposure to high light. A common link between them was found to be singlet oxygen-generated perturbations in mRNA translation that is brought about by direct singlet oxygen oxidation of RNA (Koh et al., 2023, 2021). It was shown that guanosine residues in mRNA are readily and specifically oxidized to 8-hydroxyguanosine that blocks consecutive mRNA translation. The blockage in translation releases genes that are controlled by rapidly turning over protein repressors leading to a stress transcriptome signature.

Due to its inherent instability, the oxidation damage caused by singlet oxygen occurs locally. For example, light-driven photosensitization reactions that produce singlet oxygen typically occur in the chloroplast (Flors et al., 2006; Hideg et al., 1998). Photosynthetically produced singlet oxygen was shown to damage photosynthetic machinery such as D1/2 reaction center proteins (Li et al. 2018). When singlet oxygen was produced by the photodynamic acridine orange that accumulated in the tonoplast it stimulated light-dependent vacuolar collapse and cell death. In contrast, singlet oxygen produced by rose bengal affected cytoplasmic processes (Koh et al 2016). Singlet oxygen was produced at multiple locations in the dark in many abiotic stress (Mor et al. 2014).

In the case of wounding and osmotic stress responses, a direct linkage to the activity of lipoxygenases was shown (Prasad et al. 2017; Chen and Fluhr 2018). Lipoxygenases catalyze dioxygenation of polyunsaturated fatty acids containing at least two cis-double bonds. The Arabidopsis lipoxygenase family includes six members that are categorized into types by their dioxygenation activity. Type 9 and type 13 differ in the oxygenation site of the fatty acids’ common substrate, linolenic and linoleic acid (18:8,18:2), and generate oxylipin products that show diverse biological activity (Camargo et al., 2023; Viswanath et al., 2020). Lipoxygenase type 13 is recognized for its role in jasmonic acid (JA) production (Maynard et al., 2021; Xing et al., 2019). Lipoxygenase type 9 produce other stress responses (Vellosillo et al. 2013; (Arias-Gaguancela et al., 2023; Jimenez-Aleman and Jander, 2023; Vellosillo et al., 2007).

A significant by-product of lipoxygenase activity is singlet oxygen. For example, the substrate linoleic acid, (C18:2) is oxidized to 13(S)-Hydroperoxy-9(Z),11(E)-octadecadienoic acid (13-HPOD). A minor portion of the hydroperoxide intermediates are radicalized and could spontaneously participate in bimolecular Russell reactions that resolve themselves to produce singlet oxygen (Russell, 1957). The physiological importance of this was demonstrated in mutant Arabidopsis lines with reduced lipoxygenase activity. Those lines displayed significantly less singlet oxygen production and less repression of root growth due to osmotic stress. Furthermore, supplementation of root media with linoleic acid was sufficient to stimulate singlet oxygen production. The level of lipoxygenase fatty acid substrates rapidly increased in osmotic stress, suggesting that substrate supply was a limiting component in the osmotic response (Chen et al., 2021).

At the cellular level little is known about how lipoxygenase activity and its associated singlet oxygen by-product are manifested and how the various subcellular compartments play a role. Here we show, using chemical inhibitors and mutants, that both lipoxygenase types produce singlet oxygen. By defining precise morphological stages in the subcellular response to osmotic stress we show that each lipoxygenase family has a specific contribution to cellular fate. Furthermore, during stress, the type 9 lipoxygenase was relocated from being spread evenly in the cytosol to an association with plastids. That finding was correlated with the detection of type 9 oxylipins. Additionally, small bodies producing singlet oxygen were shown to originate from the plastid and play a major role in membrane remodeling after osmotic stress. Thus, the singlet oxygen burst is correlated with cellular changes resulting in distinct morphological and physiological effects.

## Results

### Osmotic stress generates singlet oxygen in root tissue and is associated with a reduction in root elongation

Osmotic stress in roots was induced with solutions of polyethylene glycol PEG (30% W/V; -2.035 MPa) of an average of 8,000 MW (Chen and Fluhr, 2018). Short-term treatments between 10-180 min were examined to better correlate cellular changes with the appearance of singlet oxygen. In addition, to measure the effect of stress on growth, seedlings were re-plated to a standard medium, and root length was measured after 5 h. Roots exhibited a progressive reduction in growth within 10 minutes post-treatment with PEG (Figure 1 A). Singlet oxygen is short lived but can be captured by the chemical probe Singlet Oxygen Sensor Green (SOSG) (Flors et al., 2006). In contrast to singlet oxygen, the oxidized product of the SOSG reporter, oxSOSG, accumulates. After osmotic stress, it was localized mainly to cells in the elongation zone in a diverse and autonomous pattern (Figure 1 B and C). Some cells were not responsive, and others displayed dotted foci of oxSOSG scattered throughout the cell.

**Figure 1.**
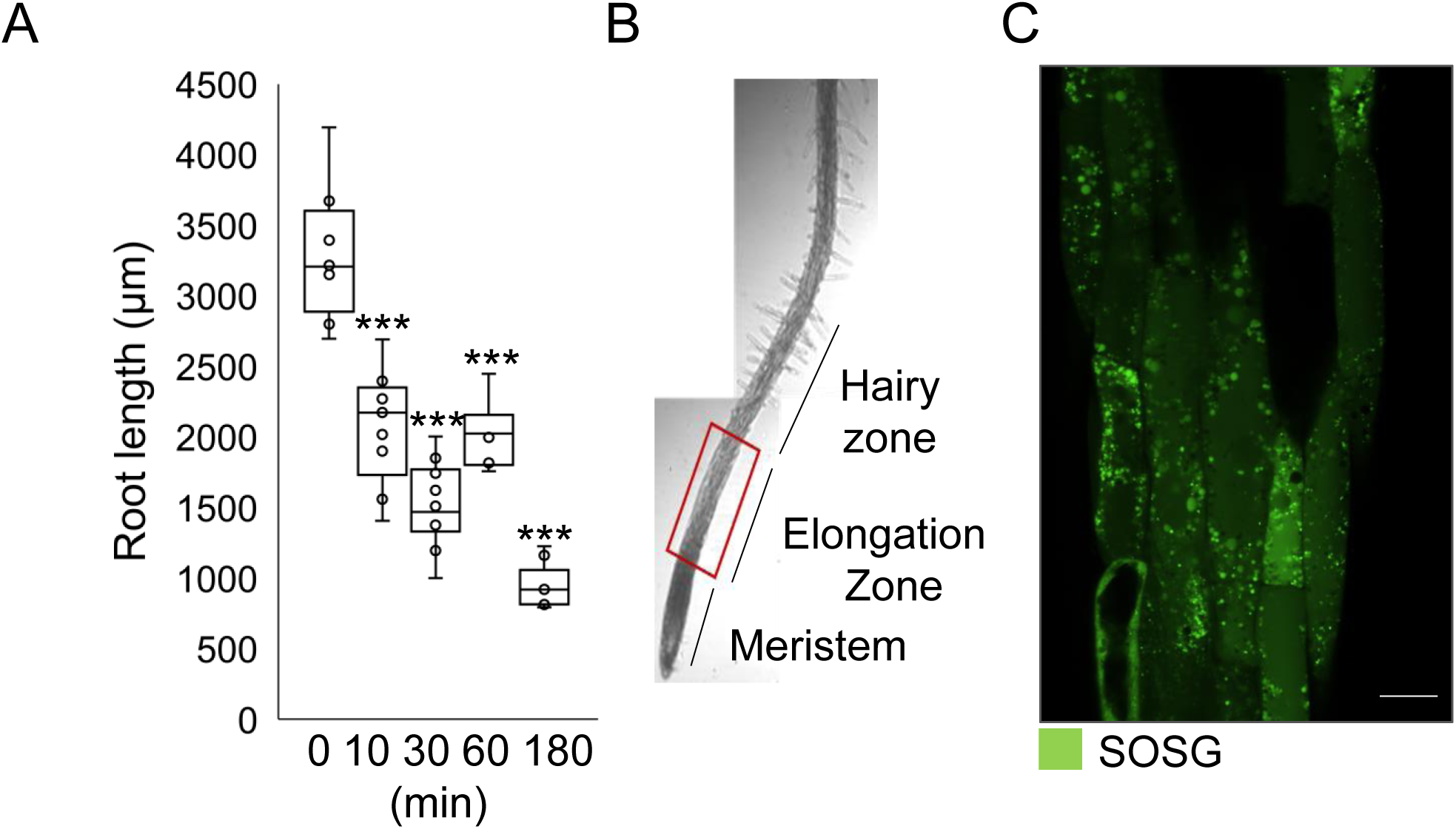
Root elongation and appearance of oxSOSG after PEG treatment. **A**, Quantification of root elongation after PEG treatment. Arabidopsis seedlings, 5 days old, pre-incubated in DDW for 30 min were treated with PEG for 0,10, 30, 60 or 180 min. The seedlings were moved to a fresh MS agar plate for 5 hours for recovery. Roots were measured under the microscope and the length quantified by ImageJ. The bars show the average root growth of 9-10 roots in treatment with SE. P value = *** < 0.001 in two-way ANOVA and Tukey test. Number of roots, n equals 46. **B**, Area of interest in the root. The red square marks the sampling area shown in (C). **C**, Visualization of oxSOSG in root cells. Arabidopsis seedlings of WT 5 days old, pre-incubated in DDW for 30 min were treated with PEG for 30 min. Seedlings were dyed with SOSG for 20 min and visualized by confocal microscopy. The layer from which examinations were made was one-third of the root width from the surface. Bar equals 20 μm.

Cells in osmotically stressed tissue were categorized, based on the accumulation of oxSOSG, by using plasmalemma and tonoplast membrane marker lines. Figure 2 A shows membrane markers in red, oxSOSG in green, and overlapping areas in yellow. Cells were categorized into 4 types (Figure 2 A and C). “N” (no response) cells exhibited singlet oxygen levels below the detection threshold. The cell and vacuole membranes in N-type cells closely resembled those in untreated root cells. Affected cells that showed oxSOSG accumulation within round foci were divided into minor response (Mi) and major response (Ma). Mi group cells featured oxSOSG-containing bodies adjacent the cell membrane and the tonoplast, with membrane marker topology like N-type cells. In cells classified under the Ma category, the tonoplast underwent segmentation and oxSOSG bodies were dispersed throughout the cytoplasm and the vacuolar compartment. In both Mi and Ma-type cells, the plasmalemma adhered to the cell wall, indicating continued cell vitality. Cells designated as “D” (dead cells) displayed a weak and uniform background of oxSOSG marker. In type D cells, the cell membrane was detached from the cell wall, enabling leakage of oxSOSG fluorescence, and the tonoplast marker was scarcely detectable.

**Figure 2.**
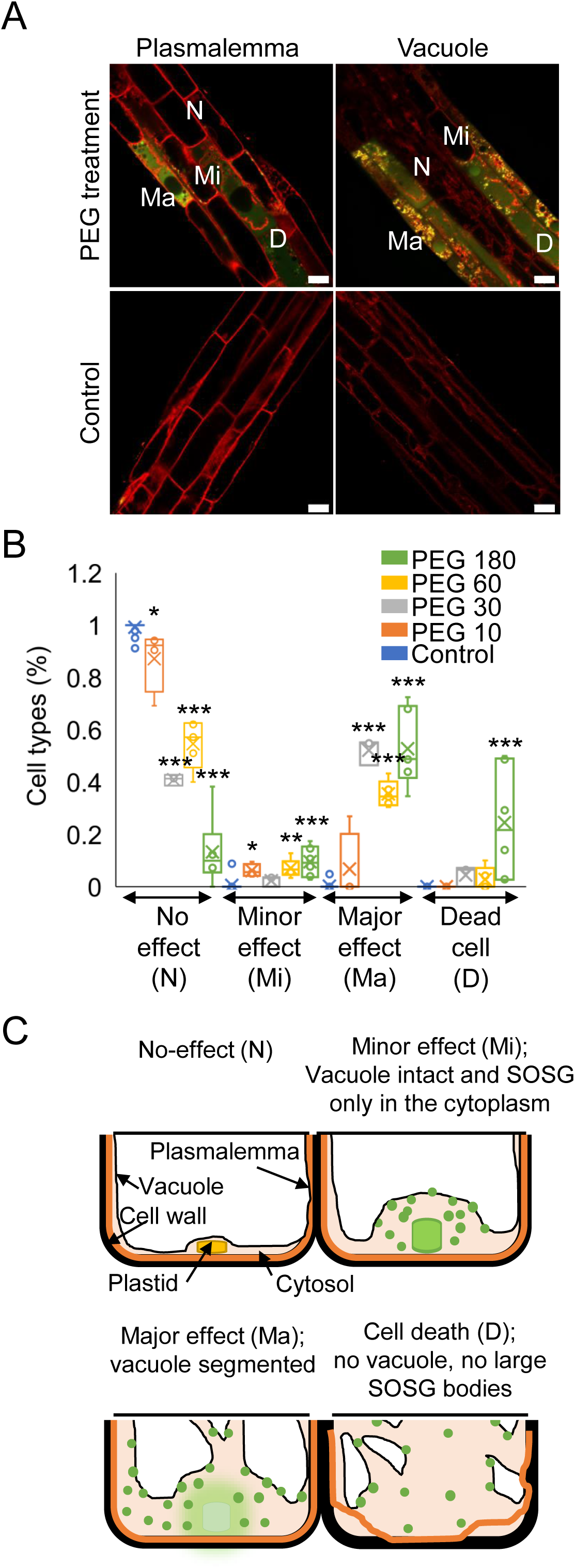
Characterization of osmotic stress by cell types. **A**, The co-localization of oxSOSG with plasmalemma, vacuolar, and plastid markers in osmotic stressed roots. Arabidopsis seedlings are transgenic lines transformed with RFP markers for, plasmalemma (pm-mcherry) and vacuole (wave 9R:VAMP711). Seedlings, 5 days old, pre-incubated in DDW for 30 min were treated with PEG for 0 and 30 min. Seedlings were dyed with SOSG for 20 min. Cell type classifications are N- no effect, Mi- minor effect, Ma- major effect, and D- dead cell. Bar equals 20 μm. **B**, Percentage of cell types after different times of stress treatment. Arabidopsis seedlings, 5 days old, pre-incubated in DDW for 30 min were treated with PEG for 0,10, 30, 60, and 180 min. Seedlings were dyed with SOSG for 20 min. The cell types were classified as in **(A)**. The cells were counted manually and divided by the total number of cells in the image. Bars show the average with SE. P value = * < 0.05, ** < 0.01, *** < 0.001 in two-way ANOVA and Tukey test. Number of roots, n equals 41. **C**, Schematic diagram of the four types of cellular responses to osmotic stress with oxSOSG localization. The scheme describes changes at the cellular level during osmotic stress. Green color represents sources of SOSG fluorescence.

Temporal examination of tissue after variable times of osmotic stress showed distinct post-osmotic stress distribution of cell types (Figure 2 B). Within 10 minutes of osmotic treatment, a detectable decrease was noted in the percentage of N-type cells compared to the control group. The percentage of N-type cells continuously declined as the duration of osmotic stress treatment increased. The population of Mi-type cells increased slightly within 10 min and remained at a constant, unchanging low level after all times, Thus, Mi likely represents a brief transitory cellular state.

In contrast, the percentage of Ma-type cells rose and remained high after 30 min. A comparison of oxSOS accumulation and cell death markers, measured by SYTOX, showed correlation (Supplementary Figure S1). Notably, while N-type cells show a positive correlation with root growth, the D-type of cell responses showed reciprocal repression of root growth (Supplementary Figure S2). Thus, the short-term cellular effects due to osmotic stress that were depicted here show a steady progression of cellular states to cell death. They align with previous results that established a correlation between singlet oxygen and cell death (Chen and Fluhr, 2018).

### Localization of singlet oxygen in oxSOSG bodies and changes in plastid morphology

Within Ma-type cells, distinct rounded structures rich in oxSOSG ranging in sizes from less than 0.1 to 10 μm² accumulated (Figure 2 A). A soluble plastid stromal marker, the FNR1 signal peptide linked to fluorescent RFP (stromal ds-RED), was used to address the origin of these bodies. N-type cells displayed plastids of conventional size and shapes (Figure 3 A, Control). However, following a 30 minute treatment, plastids swelled and became rounded (Figure 3 A, 30 min upper row). The overlap between the stromal labeling and the fluorescence of oxSOSG show that some singlet oxygen production-occurred within the root plastids (Figure 3 A, 30 min bottom row). Other, smaller rounded bodies containing oxSOSG showed no overlap with the stromal marker. Under magnification the plastids displayed protrusions of oxSOSG that did not contain stromal content (Figure 3A Mag, compare Plastid RFP, SOSG and Merge; note arrows in bottom). The observation suggests that blebbing occurs from the outer plastid membrane that does not include stromal content. The results suggest that oxSOSG accumulated in the plastid after stress and that small bodies devoid of stroma but that contain oxSOSG also originated from the plastids.

**Figure 3.**
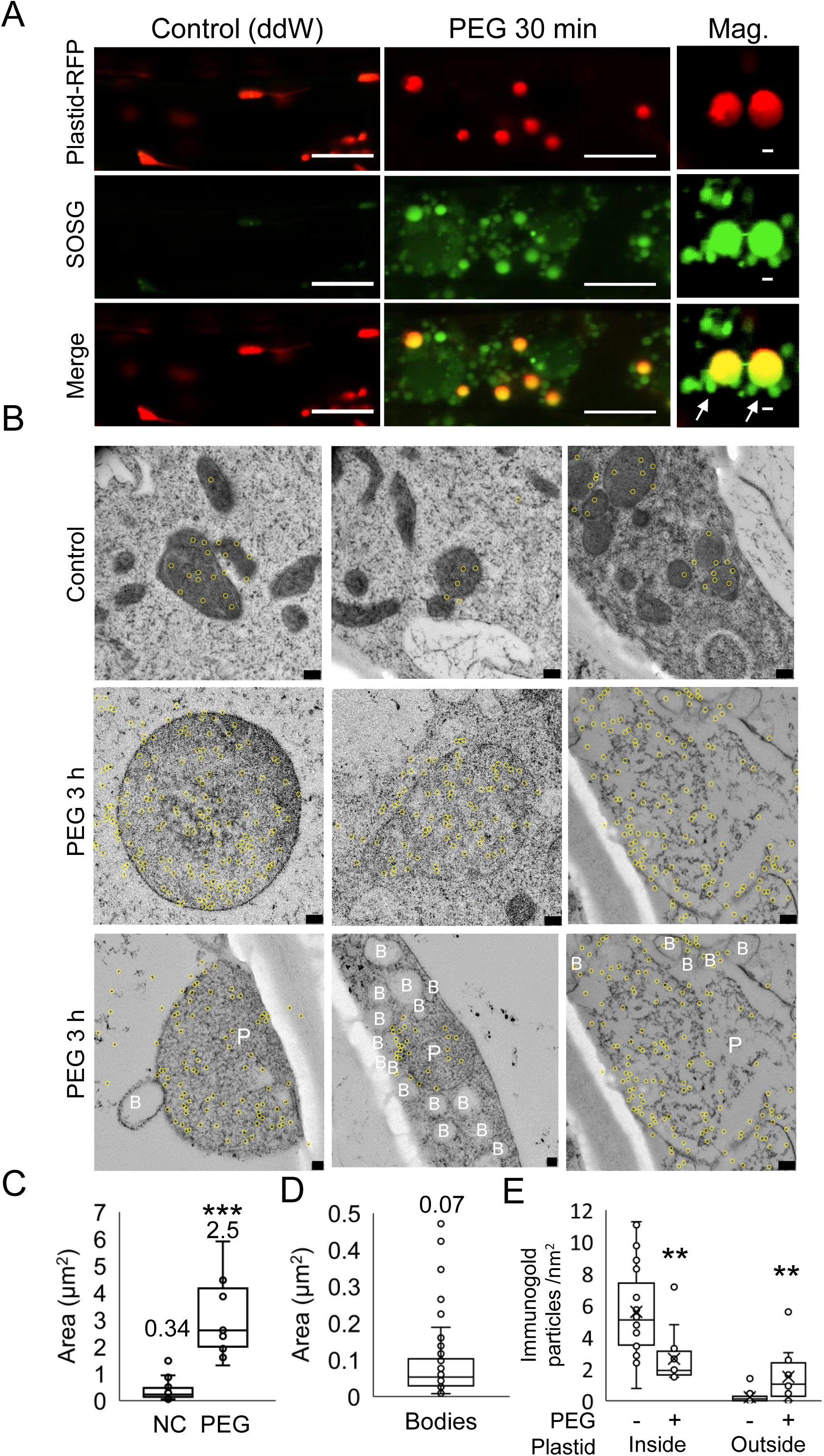
Production of singlet oxygen and changes in plastid morphology after osmotic stress. **A**, Visualization of plastids with oxSOSG co-localization by confocal microscopy. Arabidopsis seedlings expressing plastid stromal dsRED labelled Plastid-RFP, 5 days old, pre-incubated in DDW for 30 min were treated with PEG for 0- and 30-min. Seedlings were dyed with SOSG for 20 min to show oxSOG. In the control (ddW) and PEG 30 min bar equals 10 μm. In PEG Mag bar equals 1 μm. **B**, Visualization of plastids by electron microscopy. Signal peptide of the plastid stroma was marked by immunogold gold (highlighted by yellow rings). Arabidopsis seedlings of the transgenic line stromal CT-GFP, 5 days old, pre-incubated in DDW for 30 min were treated with PEG for 0- and 180-min. Samples were prepared for electron microscopy as detailed in the Materials and Methods and marked by anti-GFP gold particles. Upper row, control; Middle row treatment for 180 min. Bar equals 200 nm; Lower row, visualization of stromal free bodies that surround plastids. Plastid are labeled ‘P’ and stromal free bodies ‘B’. Bar from left to right equals 100, 100 and 500, nm. **C-E**, Quantification of EM images. Images from **(B)** were quantified using ImageJ. The bars show the average with SE. P value = * < 0.05, ** < 0.01, *** < 0.001 in t-test. Number of Cells, n equals 20. **(C)** Quantification of plastid area in μm^2^. **(D)** Distribution of area of small body size vesicles in μm^2^. **(E)** Quantification of number immunogold particles inside and outside the plastid area in nm^2^.

Higher resolution imaging of control was carried out using electron microscopy. Transgenic lines with a stromal FNR1 signal peptide fused to GFP (stromal CT-GFP) were examined with anti-GFP antibodies attached to gold particles to identify root plastids. The appearance of gold particles identified root plastids (Figure 3 B, control, gold particles are highlighted by yellow circles). Alterations in plastid morphology were evident in the cross-sections after osmotic stress and are quantified in Figure 3 C, revealing a substantial 7-fold increase in plastid size following treatment. After osmotic stress, a two-fold increase in the number of gold particles was observed outside the plastid structure compared to the control. The observation may represent stress-induced stromal leakage (Figures 3 B, middle row and Figure 3 E, quantification). Furthermore, smaller bodies appeared surrounding the plastids (Figure 3 B marked by ‘B’ in bottom row). They were measured to be of 0.07 average size (Figure 3 D, range 0.01 to 0.09 μm²). These structures lacked the stromal CT-GFP marker, similar to observations made by confocal microscopy for oxSOSG (Figures 3 A, Mag). Taken together the results suggest that the plastids undergo stressed-induced enlargement and that small bodies devoid of stroma originated from the plastids during stress.

### Response of the plastid outer membrane to osmotic stress

Confocal microscopy highlighted small stroma-free protrusions on the plastids, whilst electron microscopy showed stroma-free bodies surrounding plastids. Hence, the numerous small oxSOSG containing bodies may originate through pinching-off from the plastid outer membrane. The CRUMPLED LEAF (CRL) protein is localized to the plastid outer membrane (Asano et al., 2004; Wang et al., 2020, p. 202). To explore origins of the small oxSOSG bodies, a CRL-GFP line was subjected to osmotic stress (Wang et al., 2020). In control root cells, CRL appeared in the periphery of plastids. Some plastids contained small bodies and stromule-type structures (Figure 4 A, top row, Control, yellow arrows). After osmotic stress treatment for 30 min, plastids swelled and CRL labeling of the membrane was apparent. In addition, small extraplastidic protrusions were observed (Figure 4 A, PEG 30 min).

**Figure 4.**
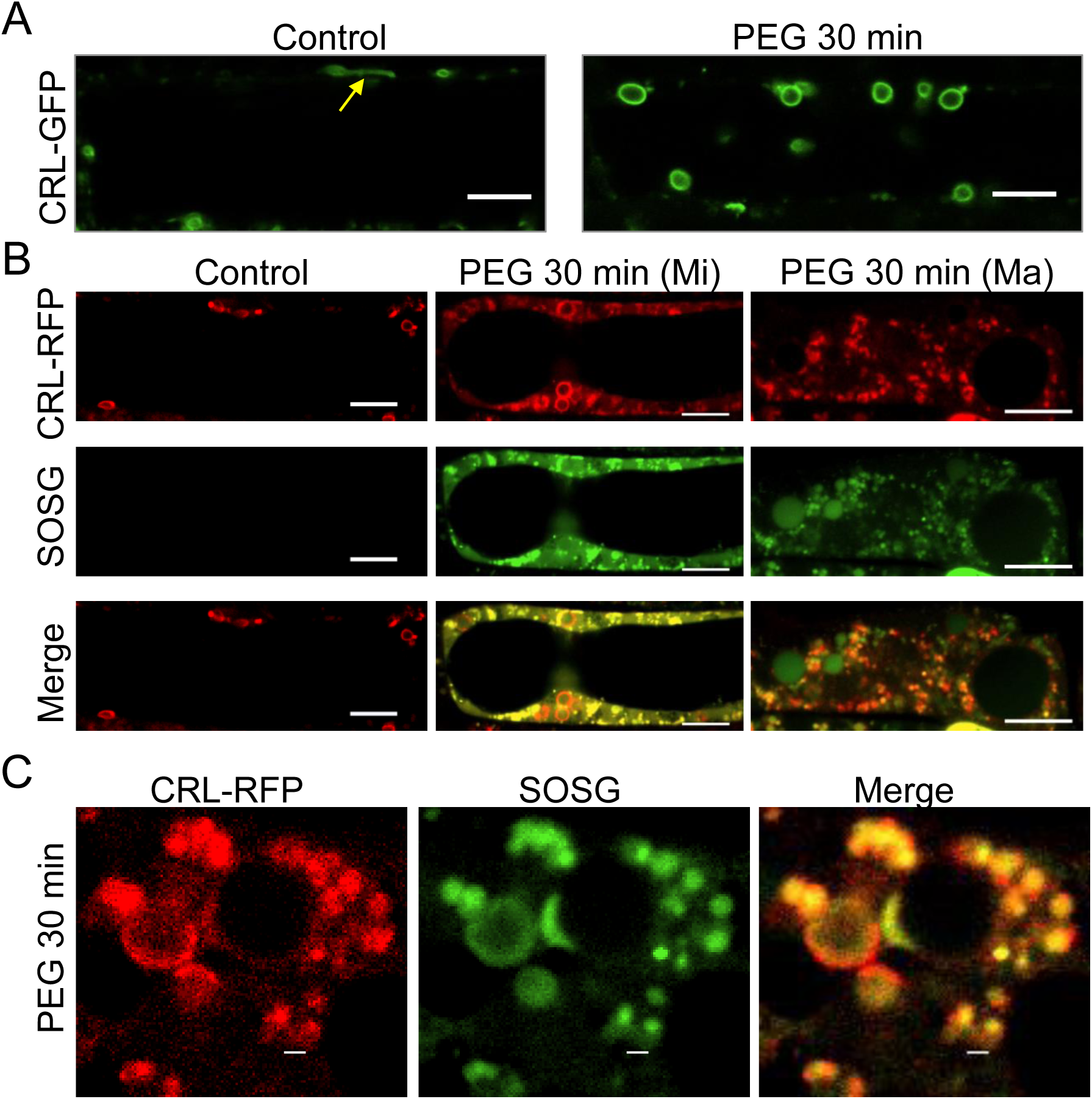
Plastid Visualization of the CRL-GFP, CRL-RFP plastid outer membrane marker and oxSOSG in osmotic stress. **A,** Plastid membrane visualization of 35S::CRL-GFP. Seedlings of CRL-GFP, 5 days old were pre-incubated in DDW for 30 min and treated with PEG for 0 or 30 min. yellow arrow shows a stromule. Bar equals 10 μm. **B,** Arabidopsis seedlings of a 35S::CRL-RFP transgenic line, 5 days old, pre-incubated in DDW for 30 min were treated with PEG for 30 min or DDW (control).The seedlings were dyed with SOSG for 20 min. Mi and Ma stage cells are shown. Bar equals 10 μm. **C,** Enlarged view as in (B) of plastid (from Ma stage cells). Bar equals 1 μm.

A CRL-RFP transgenic plant was constructed to simultaneously visualize CRL and oxSOSG markers. As shown in Figure 4 B complete overlap of CRL-RFP and oxSOSG was obtained. Within 30 min of osmotic stress blebbing of small extraplastidic CRL-RFP bodies appear, mostly emanating from the enlarged plastids (Figure 4 C). The extraplastidic bodies appear to contain oxSOSG surrounded by the CRL-RFP marker (Figure 4 C). Thus, analyses by confocal observation show that the small oxSOSG-containing bodies contain the CRL-GFP marker and likely originate from the plastid outer membrane area.

### LOX activity determines osmotic stress responses

Lipoxygenase activity was previously shown to generate singlet oxygen. Hence it is of interest to follow its impact on the cellular responses that were characterized. To this end, three modulators of lipoxygenase activity were examined. salicylhydroxamic acid (SHAM) an inhibitor of enzymatic activity; histidine (His), a scavenger of the singlet oxygen, and a µlox line in which transcripts of all LOX members have been down-regulated (Chen et al., 2021). In each case, the distribution of cellular types delineated in Figure 2 was followed after osmotic stress.

In roots, osmotic stress of 30 minutes resulted in a significant induction of oxSOSG accumulation. However, when co-treated with SHAM, His, or in the µlox line, oxSOSG was low or undetectable (Figure 5 A; Supplementary Figure S3). While the level of oxSOSG was reduced in all treatments, examination of the distribution of cell types showed interesting differences. Due to the application of SHAM, or in the µlox line, the preponderance of cells remained as N-type. In contrast, in the presence of His, a large percentage of cells were of the Ma-type, yet the percentage of D-type cells was reduced (Figure 5 B). In all cases, root growth was improved compared to osmotic stressed roots without the additional treatment (Figure 5 C). The analyses show that SHAM or the µlox background prevents the osmotic stress from evoking an initial cellular response. In contrast, His inhibits later cell death but not the initial cellular responses.

**Figure 5.**
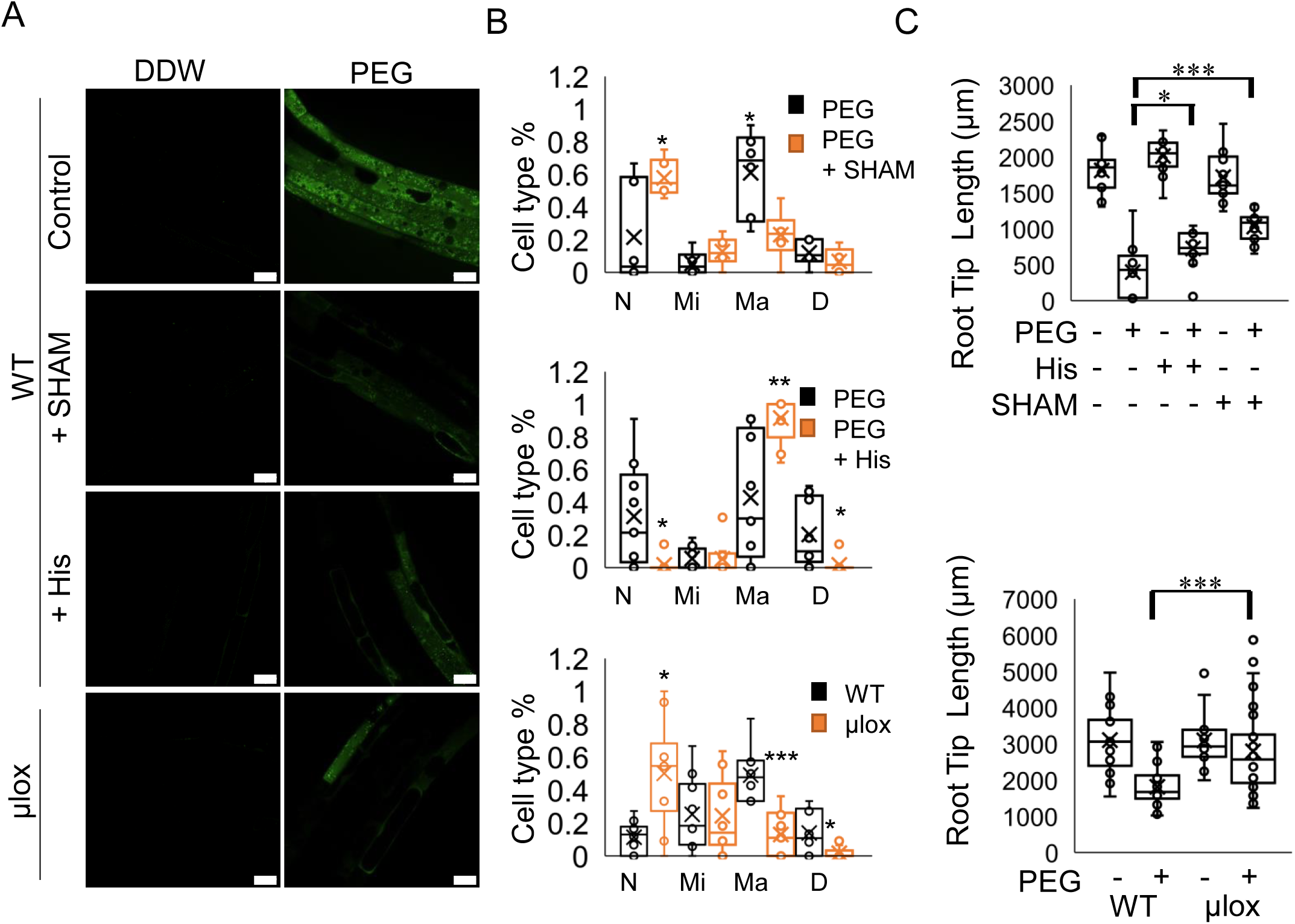
Singlet oxygen generation and root tip growth after osmotic stress in His and SHAM treated WT plants and in the µlox mutant. **A**, Visualization of the fluorescence of oxSOSG with His and SHAM and in the µlox background after osmotic stress. Seedlings were treated with PEG for 30 min in DDW, PEG, His, PEG + His, SHAM, or PEG + SHAM. Seedlings were dyed with SOSG for 20 min. Bar equals 25 μm. **B**, Percentage of cell types. Seedlings as in **(A)**. The cell types were classified as described in Figure 2. The cells were counted manually and divided by the total number of cells in the image. Bars show the average with SE. P value = * < 0.05, ** < 0.01, *** < 0.001 in student T. Test. Number of roots, n equals 48. **C**, Quantification of root elongation. Arabidopsis seedlings in the WT or µlox background, 5 days old, pre-incubated in DDW for 30 min were treated with PEG for 30 min in DDW, PEG, His, PEG + His, SHAM, or PEG + SHAM. The seedlings were transferred to a new MS agar plate for 5 hours for recovery. Seedlings were measured using ImageJ. The bars show the average growth of 14 roots in treatment with SE. P value = * < 0.05, ** < 0.01, *** < 0.001 in two-way ANOVA and Tukey test. Number of roots, n equals 80.

### LOX type 13 and type 9 motivate different responses in osmotic-stressed roots

The changes in cell distribution and root elongation using modulators of lipoxygenase activity prompt the question as to which of the two types, type 13 or type 9, or both, play a role in responding to osmotic stress. The lox2/3/4/6 quadruple mutant and the lox 1/5 double mutant retain only LOX type 9 or LOX type 13 activity, respectively. Due to osmotic stress, both lines showed reduced oxSOS accumulation, although oxSOSG reduction was more prominent in lox1/5 (Figure 6 A; Supplementary Figure S4). Interestingly, the lox1/5 double mutant, as measured by the distribution of cell types and the impact on root elongation after osmotic stress, demonstrated notable similarity to application of the SHAM inhibitor and to the results in the µlox line. Hence, lines lacking type 9 lipoxygenase activity showed a preponderance of N-type cells after osmotic stress (Figure 6 B and 6 C).

**Figure 6.**
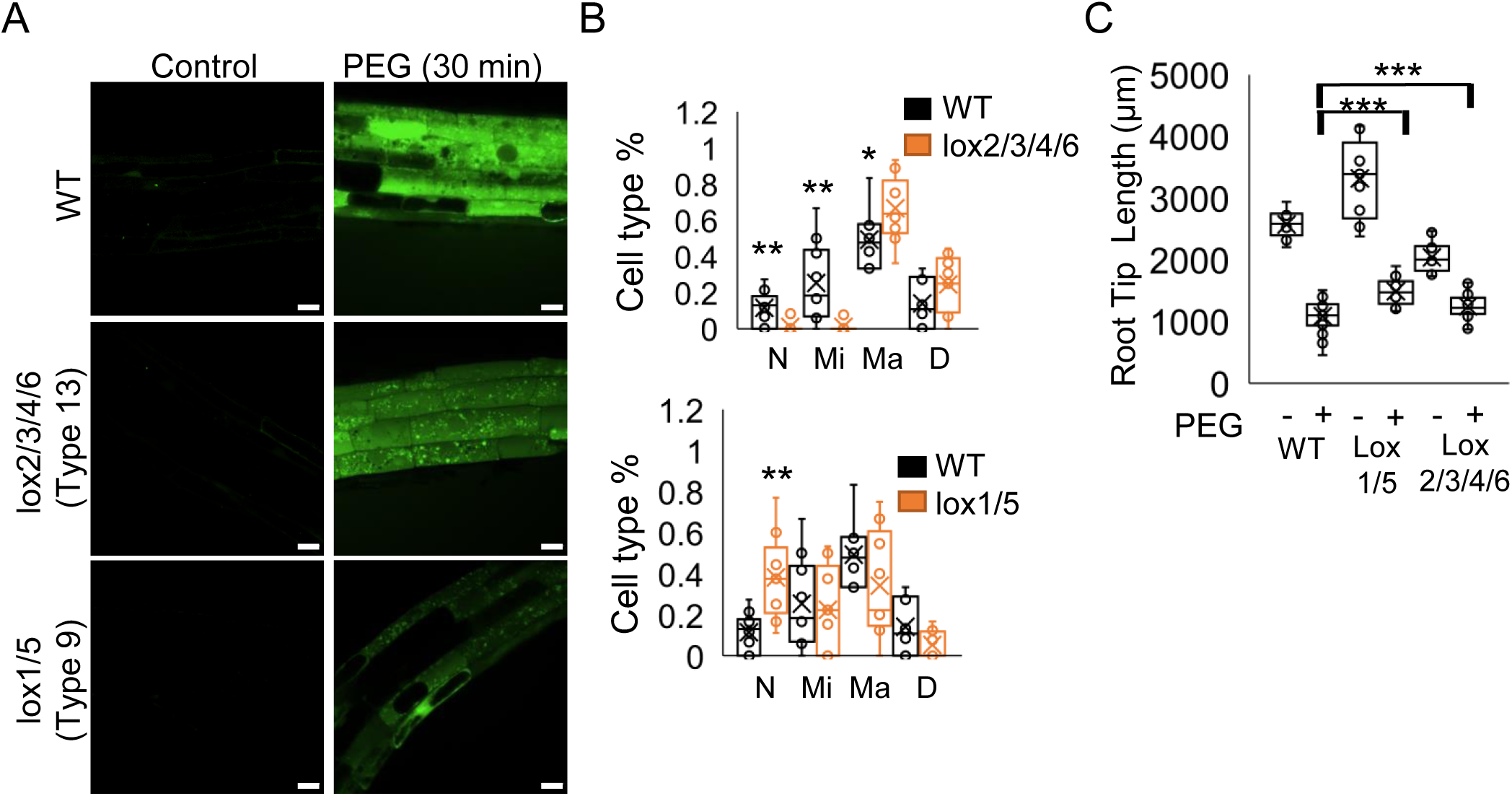
Cell types after osmotic stress in lipoxygenase mutants. **A**, Distribution of oxSOSG in osmotically stressed tissue. Arabidopsis seedlings of WT, lox2/3/4/6 quadruple mutant (lipoxygenase type 9 remains active) or lox1/5 double mutant (lipoxygenase type 13 remains active). Seedlings, 5 days old, pre-incubated in DDW for 30 min. were treated with PEG for 30 min or DDW (control). Seedlings were dyed with SOSG for 20 min. Bar equals 20 μm. **B**, Percentage of cell type. Seedlings as in **(A)**. The cell types were classified as described in Figure 2.: The cells were counted manually and divided by the total number of cells in the image. Bars show the average with SE. P value = * < 0.05, ** < 0.01, *** < 0.001 in student T-test. Number of roots, n equals 27. **C**, Quantification of root elongation. Arabidopsis seedlings of WT, lox2/3/4/6 or lox1/5, 5 days old, pre-incubated in DDW for 30 min and were treated for 30 min in DDW or PEG. The seedlings were transferred to a new MS agar plate for 5 hours for recovery. Seedlings were measured by ImageJ. The bars show the average root growth of 14 roots in treatment with SE. P value = * < 0.05, ** < 0.01, *** < 0.001 in two-way ANOVA and Tukey test. Number of roots, n equals 72.

In contrast, the lox2/3/4/6 quadruple mutant that retains only lipoxygenase type 9 activity resembles His treatment, where cells progress to Ma-type level of damage but do not die (Figure 6 B). In both mutant classes, a notable improvement in root elongation after osmotic stress was realized (Figure 6 C). Thus, cell death and the correlated root growth are mitigated by either type of LOX activity but in a different manner.

### Changing patterns of LOX localization during osmotic stress

To assess a potential connection between the differential effect of members of the lipoxygenase family and their cellular distribution, lines of LOX3-RFP (representing type 13 activity) and LOX5-RFP (representing type 9 activity) were analyzed concomitantly with oxSOSG accumulation. In non-stressed control roots, LOX3 accumulated within plastids (Figure 7 A, control). In Mi and Ma-type stages of osmotic stressed roots, LOX3 and oxSOSG fluorescence overlapped in the plastids (Figure 7 A, Mi and Ma). Interestingly, in the Ma -type cells, protrusions of oxSOSG and LOX3-RFP appear and overlap. This is in contrast to the stromal dsRED that was not found in the protrusions, although oxSOSG was found (compare Figure 7 A, Mag, bottom row and Figure 3 A).

**Figure 7.**
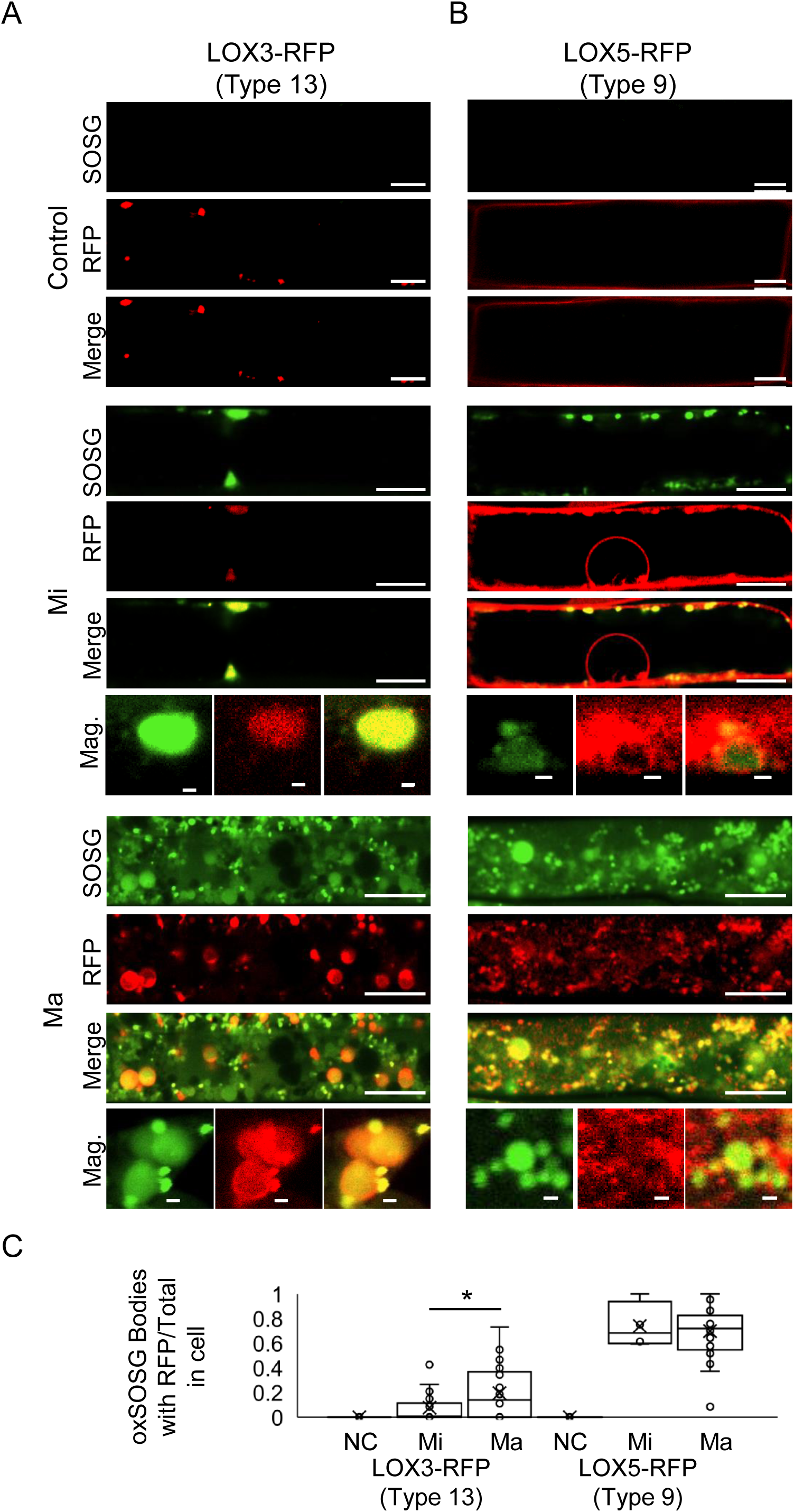
Localization of LOX5-RFP, LOX3-RFP and oxSOSG in osmotic stress. **A-B**, Arabidopsis seedlings of 35S::LOX3-RFP **(A)** and 35S::LOX5-RFP **(B)** transgenic line, 5 days old, pre-incubated in DDW for 30 min were treated with PEG for 30 min or DDW (control). Seedlings were dyed with SOSG for 20 min. images presenting control cell (Control), minor affected cell (Mi) and major affected cell (Ma), Bar equals 10 μm. The row in Mi and Ma marked Mag is a magnification of plastid), Bar equals 1 μm. **C**, Quantification of co-localization of oxSOSG with LOX5-RFP and LOX3-RFP. Images as described in **(A-B)** were measured for the signals of oxSOSG and RFP using ImageJ. The number of bodies where oxSOSG and LO3/5-RFP overlapped were divided by the total number of oxSOSG bodies. That number was then normalized by the cell area for each experiment. Control (NC), minor affected cell (Mi) and major affected cell (Ma). Bars show the average with SE. P value = * < 0.05, ** < 0.01, *** < 0.001 in student T-test. Number of Cells, n equals 200.

In contrast, LOX5-RFP accumulated evenly in the cytosol and did not appear to interact particularly with plastids (Figure 7 B, control). However, in Mi-type cells, LOX5 displayed a more punctate appearance in the cytoplasm and appear to overlap with plastids (Figure 7 B, Mi, merge). Although higher magnification microscopy shows that it surrounds the plastid (Figure 7 B, Mi, Mag). To further resolve this localization, a double transgenic line with the plastid stroma marker (stromal CT-GFP) and LOX5-RFP was examined and demonstrated that LOX5-RFP is in close proximity of the plastid in Mi-type cells (Figure 8 A). In Ma-type cells, after 30 min of osmotic stress a degree of overlap between LOX5 and numerous oxSOSG bodies was observed (Figure 7 B, Ma, Merge). In contrast, the overlap of oxSOSG bodies with LOX3-RFP was less extensive (compare Figure 7 A and 7 B, Ma, Merge and quantified in 7 C).

**Figure 8.**
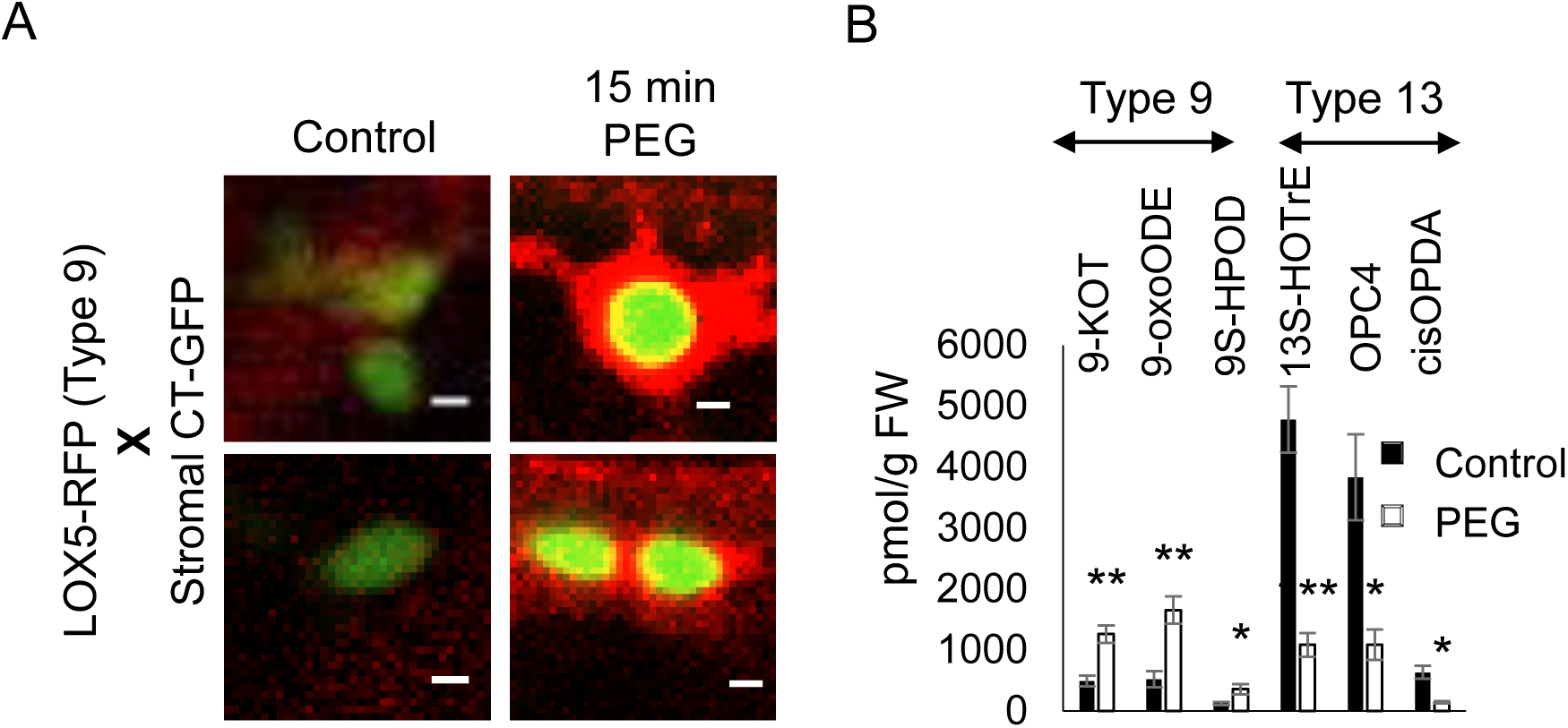
Localization of LOX5-RFP and metabolic activity associated with lipoxygenase type 9 and type 13 in osmotic stress. **A,** Co-localization of stromal CT-GFP with LOX5-RFP. Arabidopsis seedlings of LOX5-RFP/stromal CT-GFP double transgenic line, 5 days old, pre-incubated in DDW for 30 min were treated with PEG for 0 and 15 min. Bar equals 1 μm. **B,** Quantification of oxylipins that are indicative of type 13 and type 9 lipoxygenase activity. Arabidopsis seedlings, 5 days old, pre-incubated in DDW for 30 min were treated with PEG for 0 and 3 h. Roots were cut and immediately transferred to liquid nitrogen. The oxylipins were extracted and analyzed by mass spectrometry as described in the Methods. Bars are the fold change of the treatment divided by the control level with SE. P value = * < 0.05, ** < 0.01, *** < 0.001 in two-way ANOVA and Tukey test. Number of samples, n equals 16. Every group has 4 technical repeats. The concentration (nmol/g FW) was calculated using internal standards.

Lipoxygenase type 9 was juxtaposed to small oxSOSG bodies (Figure 7 B). In addition, mutants which lack type 9 lipoxygenase activity were highly reduced in the accumulation oxSOSG (Figure 6). To examine the possible ramifications of these observations, the downstream products of LOX enzymatic activity were measured. Three major oxylipin products were quantified after osmotic stress. As shown in Figure 8 B, the main type 9 detectable products of LOX activity were 9-KOT, 9-oxoODE, and 9S-HPOD. All increased after osmotic stress. In contrast, lipoxygenase type 13 products, 13S-HOTrE, OPC4, and cisOPDA significantly decreased from their basal control level after osmotic stress. The preferential detection of type 9 metabolic products is consistent with the overlap between lipoxygenase type 9 and oxSOSG accumulation.

## Discussion

Here we report the cellular manifestations of osmotic stress in the root through the lens of singlet oxygen production. Most studies about singlet oxygen are confined to aerial organs where the creation of singlet oxygen through photodynamic processes occurs in high light or in mutant plants (Mattila et al., 2023; Tano et al., 2023; Zhang et al., 2023). In those cases, dysfunction of photosynthesis and the production of singlet oxygen generated chemical messengers were shown to dramatically effect plant physiology (Li and Kim, 2023). Yet, in addition to photodynamic pathways, singlet oxygen can be produced in measurable amounts by lipoxygenases and other enzymatic activity in plants and animals (Chen et al., 2021; Murotomi et al., 2023; Prasad et al., 2017). However, the biological ramifications of such production were obscure and are advanced here.

As shown here osmotic stress stimulated profound cellular and organelle morphological changes, induced singlet oxygen production, fueled cell death and impacted on root growth. These changes represent ordered stress-induced progressions. The early occurrence of singlet oxygen was traced to plastids, but the subsequent and more major manifestation occurred in newly formed oxSOSG bodies. The latter bodies appeared to bud-off from the plastids. Importantly, the sequential process of cellular damage and final cell death incurred by osmotic stress was made rescindable, either by reducing the activity of lipoxygenases, or by scavenging for its by-product, singlet oxygen. Thus, mutants of specific lipoxygenase or application of its inhibitors prevented cells from entering the catastrophic sequence that leads to cell death. In contrast, the reduction of the oxidation potential of singlet oxygen without reduction in enzymatic activity, did not affect the changes in cellular morphology but did prevent cell death.

Stress has been shown to affect plastid morphology by causing loss of integrity, swelling and loss of fluidity in shape (Kwon et al., 2013; Wilson et al., 2014). The Mechanosensitive Channel of Small Conductance-Like2 (MSL2) and MSL3 mutants showed hallmarks of drought or environmental osmotic stress that included rounded enlarged plastids and constitutive induction of osmotic responses. Swelling in plastids could be due to accumulation of osmolytes induced by the high-water potential, and once roots were returned to lower potentials the plastids swell.

Strikingly, 60% of the cells of osmotic stressed roots showed no effect in µlox and SHAM treated plants whereas 12-20% of the cells showed no effect in PEG treated control roots (Figure 5 B).This results suggest that lipoxygenase catalytic activity either affects remodeling of plastid membrane structure or produces signals released by the downstream oxylipin products of lipoxygenase (Cook et al., 2021). Interestingly, a type 13 lipoxygenase (LOX2) was shown to be responsible for chloroplast breakdown during senescence in Arabidopsis (Springer et al., 2016). In that case, the lipoxygenase was traced to the chloroplast and its envelope. The authors hypothesized that lipoxygenase activity could introduce holes into the plastid envelope stimulating substantial release of stromal proteins that contributes to chloroplast breakdown and autophagy during senescence. Similarly, LOX3-RFP, a type 13 lipoxygenase, could clearly be traced to protrusions in the plastid membrane (Figure 7 A, bottom). In addition, changes in genes involved in the jasmonic acid pathway and massive plastid lipid remodeling were reported in barley flag-leaf senescence that could involve lipoxygenase activity (Cohen et al., 2022). Such scenario is consistent with lipoxygenase-dependent osmotic stress-induced plastid breakdown shown here.

Work here with multiple knockouts and fluorescently tagged lipoxygenases further refined the interaction of these enzymes with the root plastids during osmotic stress. Based on the distribution of cell types, the absence of type 9 activity after osmotic stress resembled application of the inhibitor SHAM or observations made in the µlox line. This would implicate type 9 lipoxygenases as playing a key role in plastid remodeling and initiating cellular breakdown. In contrast, the absence of type 13 lipoxygenases resembled the cellular distribution that occurs after scavenging of singlet oxygen i.e. preventing cell death but not the progression to cellular disruption. While lipoxygenases of the type 13 have been localized to chloroplasts (Springer et al., 2016) the type 9 have more diverse localizations that include plastids, cytoplasmic and tonoplast (Tolley et al., 2018).

Remarkably during osmotic stress, the type 9 lipoxygenase, LOX5, was mobilized from a uniform cytoplasmic distribution to one clumped with plastids and oxSOSG bodies (Figures 7 B, 8 A). The association of lipoxygenase type 9 with the chloroplast and extraplastidic small bodies impacted on the type of oxylipins that accumulated. Oxylipins that originated from type 9 activity increased while those indicative of type 13 activity decreased. Other tissue or plant species may respond differently to osmotic stress. For example, in Arabidopsis foliar tissue, dehydration stress transiently induced the accumulation of OPDA that is the product of type 13 activity (de Ollas and Dodd, 2016; Savchenko et al., 2014). In rice, transient increases in JA could be detected after ½ h in roots (Tang et al., 2020). Thus, measurements of metabolic activity may show time and species dependence.

Oxylipins, the results of type 9 lipoxygenase activity are of special interest as they play defined roles in plant defense. 9-hydroxyoctadecatrienoic acid was shown to control root development and plant defense by activating the Brassinosteroid pathway that signal for cell wall-based responses (Marcos et al., 2015; Vellosillo et al., 2007). In Arabidopsis roots, 9-HOT induces a characteristic waving phenotype, accompanied by oxidative stress and callose deposition (Marcos et al., 2015; Vellosillo et al., 2007). Paradoxically, the lack of the type 9 LOX5 inhibited Green Peach Aphid colonization (Nalam et al., 2012). This was explained by the finding that the presence of LOX5 increases water content in the vascular tissue which is conducive to aphid growth.

Based on the intense fluorescent signal emanating from oxSOSG, the small oxSOSG bodies are apparent hotbeds of lipoxygenase activity. Several observations suggest that oxSOSG bodies originate from the plastid. Using multiple fluorescent markers, and observation by EM, budding vesicles that do not contain stromal material was observed in the plastids after osmotic stress (Figure 3 A, 3 B and 7 A). The buds contain the CRUMPLED LEAF (CRL) outer membrane envelope marker (Figure 4 A) but do not contain stroma, which differentiates them from stromules (Delfosse et al., 2018). Budding-off, a process where smaller plastids separate from the main plastid body, has been observed under specific circumstances, particularly in mutants with disrupted division ring localization (Pyke, 2013). It was hypothesized that CRL plays a role in plastid division and the distribution of plastids into daughter cells (Sugita et al., 2012; Wang et al., 2020).

The extraplastidic formation of small bodies reported here is produced from the plastid outer membrane without containing stromal material. It thus differs from small bodies of chloroplast origin that were described for senescence. In dark stressed green tissue, an autophagosomal-dependent process produces small Rubisco-containing bodies that are pinched off from the chloroplast (Ishida et al., 2008). Additionally, there exists an autophagy-independent chloroplast vesiculation pathway that produces vesicles within the chloroplast, as well as extraplastidic bodies containing both stromal and thylakoid membrane proteins (Wang and Blumwald, 2014) In both cases the bodies are targeted to the vacuoles for turnover of their contents to ameliorate starvation conditions (Barros et al., 2023). The extraplastidic bodies shown here may merge with the vacuole as shown in Figure 2 A and are thus involved in membrane turnover.

One scenario suggests that osmotic stress causes a separation of the plastid double membrane where a bud would contain material from the periplasmic space. Indeed, hypertonic conditions cause the inner envelope membrane to separate from the outer membrane and are used in the purification of separated inner and outer envelope membranes (Cline et al., 1981). This possibility is of interest, as such a bud would contain remnants of the periplasmic space (i.e material between the inner and outer membranes). The plastid supplies acyl-CoA moieties to the ER where membrane components are first hydrolyzed into free fatty acids (FFA) through the Fatty acid Export 1/Long-chain acyl-CoA synthesis (FAX1/LACS) mechanism situated in the inner membrane (Wang et al., 2020). Thus, buds composed of the outer membrane that also contain periplasmic material would provide a rich source of substrates for the type 9 lipoxygenases that surround them. Singlet oxygen is produced by Russel reactions that require a sufficient local bimolecular concentration of the hydroperoxide product of lipoxygenase activity (Russell, 1957). Small vesicles rich in substrate and surrounded by lipoxygenases would provide an ideal base for such reactions. In addition, it would provide a mobile platform whereby the very short-lived ROS can travel long distances in the cell.

What sort of signal may singlet oxygen provide? As a by-product of a lipoxygenase reaction, singlet oxygen minimally serves as an indirect reporter of enzymatic activity. That activity is shown here to be essential for enabling the transition to cell death in osmotic stress. However, it likely serves a direct functional role in signaling cell death since the scavenging of singlet oxygen impacts on cell fate by preventing cell death and improving the chances for recovery. Cell death by singlet oxygen can occur through direct inactivation of the membrane, protein and nucleic acid components of the cellular machinery (Koh et al., 2021; Koh and Fluhr, 2016). Furthermore, singlet oxygen can modify the oxylipins produced and change their potency (Izquierdo et al., 2021). Importantly, lipoxygenase activity does not have to be induced by the stress as their basal activity is chiefly limited by the availability of fatty acid substrate (Chen et al., 2021). This accounts for the rapidity of the cellular response and would mean that the activation of lipases that provide substrate to lipoxygenase are a rate-limiting factor in osmotic stress responses and should be investigated for this function.

## Materials and Methods

### Plant growth conditions, stress treatments and elongation assay

*Arabidopsis thaliana* (ecotype Columbia-0) seedlings (5 to 7 d old) were grown under white light in a 16-h-light/8-h-dark cycle at 21°C on Murashige and Skoog medium, supplemented with 1% (w/v) Sucrose and 0.8% (w/v) Phytoagar (Invitrogen). For root elongation assay, seedlings, were pre-incubated in DDW for 30 min and then removed to a new MS agar plate for 5 h of recovery. The roots were then imaged using Prime BSI (photometrics) camera and measured using ImageJ. For osmotic stress treatment, polyethylene glycol 8,000 MW was used (Sigma-Aldrich). For PEG solutions, 30% (w/v) of PEG was dissolved in double distilled DDW. Where indicated solutions were supplemented with Histidine (His) 5 mM or salicylhydroxamic acid (SHAM) 10 mM.

### Plant lines

The plant lines, 35S::LOX3-RFP, 35S::LOX5-RFP, 35S:: PGL34-RFP, CRL::RFP were full-length cDNA cloned by the golden braid method using the GB omega plastid in Sapp. 9 (Sarrion-Perdigones et al., 2013). The µlox line was from Chen et al. 2021. The lox1/5 double mutant was derived by crossing from the tDNA insertion mutants SALK_012188 and SALK_050933. The quadruple mutant lox2/3/4/6 was generously given by Edward E. Farmer (Chauvin et al., 2013). Transgenic markers: plasmalemma marker (PM-mCherry), vacuole (wave 9R:VAMP711), plastid (FNR1 signal peptide ; stromal ds-RED and stromal CT-GFP), pre-vacuolar bodies (wave 7R: RabF2a (Rha1) are from the Niko Geldner Lab (Geldner et al., 2009; Nelson et al., 2007). CRL-GFP was generously given by Chanhong Kim (Wang et al., 2020).

### Confocal microscopy, stains, and image analysis

All images were taken with a model A1 Nikon Confocal Microscope with a GaAsp detector. In each figure, the images displayed are of equal laser power and acquisition settings unless indicated. SOSG was from Thermo Fisher Scientific. Staining was performed by incubation with 100 µM SOSG diluted in DDW in the dark for 20-30 min and washed three times. Propidium iodide (PI) stock was 15 µM. SYTOX FR stock 5 mM Solution in DMSO from Thermo Fisher Scientific. Excitation of SOSG and GFP lines was at 489 nm, and emission was at 525 nm. For PI and RFP excitation was in 561 nm and emission at 593 nm. Excitation of SYTOX FR 638 nm and emission at 700 nm. Confocal image analyses were performed using ImageJ.

### Electronic microscopy

Arabidopsis roots were excised, immersed in 1-Hexadecene as cryoprotectant and immediately fixed by high pressure freezing (HPF) using the Leica EM ICE (Leica Microsystems GmbH, Germany). Freeze substitution and embedding of the HPF-fixed samples were conducted in a temperature-controlled device, AFS2 (Leica Microsystems GmbH, Germany) at − 90 °C for 55 h, using 0.1% Uranyl Acetate in dry acetone. The temperature was then raised to −45°C (5 °C/h) for 9 h followed by three acetone washes. Infiltration with Lowicryl HM20 (Electron Microscopy Sciences, USA) was conducted at increasing concentrations (10%, 25%, for 2 h each). The temperature was then raised to −25 °C (5 °C/h) and infiltration with higher concentrations of Lowicryl HM20 (50%, 75%, 2 h each) was conducted. Finally, 100% Lowicryl HM20 was exchanged three times for every 10 h followed by polymerization under UV light for 48 h. The temperature was increased to 20 °C (5 °C/h) and left under UV light for 48 h. Ultrathin sections were obtained using an EMUC7 ultramicrotome (Leica microsystems) and were mounted on formvar coated 200 mesh nickel grids. Sections were stained with Reynolds lead citrate and digital electron micrographs were acquired using a Thermo Fisher Scientific Tecnai T12 transmission electron microscope operating at 120 kV and equipped with a bottom mounted TVIPS TemCam-XF416 4k x 4k CMOS camera.

### Lipids extraction identification quantification and analysis

Frozen samples (10-30 mg; fresh weight) were ground to fine powder with Zirconium balls and 1 ml cold (−20°C) methanol/water/formic acid (15/4/1 v/v/v) was added with 25 µl of 0.08 ug/ml of each one of the marker oxylipins d4-9-HODE, d4- 12 13-DiHOME, d4- 9 10-DiHOME, d6-20-HETE, d8-12S-HETE, d8-5S_HETE, d4-LTB4, d4-15-deoxy-Δ12 14-PGJ2, d11-14,15-diHET, d4-PGD2, d4-PGF2α, d4-6-keto PGF1α, d4-TXB2, d5-LTE4 and d4-PGE2 as stable isotope-labeled internal standards (IS). The samples were sonicated 3 times x 10 min, ground again, centrifuged 10 min at 14000 rpm and 50 µl transferred to vials for Oxylipins analysis. Oxylipins were measured using UPLC ACQUITY (Waters Corp., MA, USA) system coupled to a TQ-XS mass spectrometer (Waters Corp., MA, USA). The chromatographic separation was performed on an ACQUITY UPLC BEH C18 column (2.1×100 mm, i.d., 1.7 μm) (Waters Corp., MA, USA). Mobile phase A consisted of 3% acetonitrile in DDW when mobile phase B was 100% acetonitrile ; both supplemented with 0.1% acetic acid. The column was maintained at 35°C; flow rate of mobile phase was 0.3 ml/min. The measurement was performed with two Multiple reaction monitoring traces for each compound. Capillary voltage was 2.5 kV. Data were processed with MassLynx software with Targetlynx (Waters). Quantification of 9-KOT, 9oxo-ODE, 13S-HOTrE, 9S-HOTrE, JA, OPC4, cisOPDA, JA-Ile and IAA was done against external calibration curves, using analyte/internal standard peak ratio.

## Acknowledgments

This research was supported by a grant from the ISRAEL SCIENCE FOUNDATION; Grant No. 2106/21. We thank Dr. Edward E. Farmer for the quadruple lox2/3/4/6 mutant. We thank the Dr. Niko Geldner for the generous gift of the WAVE lines. We thank Dr. Chanhong Kim for kindly providing us with the CRL-GFP line. Special thanks to Dr. Yoseph Addadi and Dr. Ofra Golani from ‘MICC cell observatory’ unit of the Life Sciences Core Facilities in the Weizmann Institute for consulting in the field of microscopy, and to Mrs. Adi Havosha and Dr. Irina Panizel for their dedicated technical work. Electron microscopy studies were conducted at the Irving and Cherna Moskowitz Center for Nano and Bio-Nano Imaging at the Weizmann Institute of Science.

## Author Contributions

D.C.H.: conceptualization, methodology, investigation, software, formal analysis, and writing-original draft, review, and editing. T.C.: conceptualization, designed and created the transgenic lines of LOX3-RFP and LOX5-RFP constructs. L.S.: carried out the root elongation experiments. N.D.: carried out all the electron microscope preparation and sampling. M.I.: carried out all the mass spectrometry and their analysis supervised by S.M. G.F.: sampled all the spectral imaging. R.F: conceptualization, supervision, writing-original draft, review, and editing.

## Data Availability Statement

Data supporting the findings of this work are provided in the main text and the supporting information files. All data and materials used in this study will be available from the corresponding author.

## Figure legends of the Supplemental data

**Supplementary Figure S1.** Relative area of oxSOSG and number of dead cells in osmotic stressed roots.

**Supplementary Figure S2.** Correlation between cell type after osmotic stress and root elongation.

**Supplementary Figure S3**. Intensity of oxSOSG in small bodies of Ma-type cells.

**Supplementary Figure S4.** Visualization and quantification of singlet oxygen and cell death in µlox, LOX 13- and type 9 background mutants.

## Abbreviations

LOX: lipoxygenase
CRL: *CRUMPLED LEAF*
PEG: polyethylene glycol
His: histidine
SHAM: salicylhydroxamic acid
FFA: free fatty acids.

## References

Arias-Gaguancela, O., Aziz, M., Chapman, K.D., 2023. Fatty acid amide hydrolase and 9-lipoxygenase modulate cotton seedling growth by ethanolamide oxylipin levels. Plant Physiol. 191, 1234–1253. 10.1093/plphys/kiac556

Asano, T., Yoshioka, Y., Kurei, S., Sakamoto, W., Sodmergen, Machida, Y., 2004. A mutation of the CRUMPLED LEAF gene that encodes a protein localized in the outer envelope membrane of plastids affects the pattern of cell division, cell differentiation, and plastid division in Arabidopsis. Plant J. 38, 448–459. 10.1111/j.1365-313X.2004.02057.x

Barros, J.A.S., Cavalcanti, J.H.F., Pimentel, K.G., Magen, S., Soroka, Y., Weiss, S., Medeiros, D.B., Nunes-Nesi, A., Fernie, A.R., Avin-Wittenberg, T., Araújo, W.L., 2023. The interplay between autophagy and chloroplast vesiculation pathways under dark-induced senescence. Plant Cell Environ. 46, 3721–3736. 10.1111/pce.14701

Ben Rejeb, K., Lefebvre-De Vos, D., Le Disquet, I., Leprince, A.-S., Bordenave, M., Maldiney, R., Jdey, A., Abdelly, C., Savouré, A., 2015. Hydrogen peroxide produced by NADPH oxidases increases proline accumulation during salt or mannitol stress in Arabidopsis thaliana. New Phytol. 208, 1138–1148. 10.1111/nph.13550

Blum, A., 2017. Osmotic adjustment is a prime drought stress adaptive engine in support of plant production. Plant Cell Environ. 40, 4–10. 10.1111/pce.12800

Camargo, P.O., Calzado, N.F., Budzinski, I.G.F., Domingues, D.S., 2023. Genome-wide analysis of lipoxygenase (LOX) genes in angiosperms. Plants. 12, 398. 10.3390/plants12020398

Chauvin, A., Caldelari, D., Wolfender, J.-L., Farmer, E.E., 2013. Four 13-lipoxygenases contribute to rapid jasmonate synthesis in wounded *Arabidopsis thaliana* leaves: a role for lipoxygenase 6 in responses to long-distance wound signals. New Phytol. 197, 566–575. 10.1111/nph.12029

Chen, T., Cohen, D., Itkin, M., Malitsky, S., Fluhr, R., 2021. Lipoxygenase functions in singlet oxygen production during root responses to osmotic stress. Plant Physiol. 185, 1638–1651. 10.1093/plphys/kiab025

Chen, T., Fluhr, R., 2018. Singlet oxygen plays an essential role in the root’s response to osmotic stress. Plant Physiol. 177, 1717–1727. 10.1104/pp.18.00634

Cline, K., Andrews, J., Mersey, B., Newcomb, E.H., Keegstra, K., 1981. Separation and characterization of inner and outer envelope membranes of pea chloroplasts. Proc. Natl. Acad. Sci. U. S. A. 78, 3595–3599. 10.1073/pnas.78.6.3595

Cohen, M., Hertweck, K., Itkin, M., Malitsky, S., Dassa, B., Fischer, A.M., Fluhr, R., 2022. Enhanced proteostasis, lipid remodeling, and nitrogen remobilization define barley flag leaf senescence. J. Exp. Bot. 73, 6816–6837. 10.1093/jxb/erac329

Cook, R., Lupette, J., Benning, C., 2021. The role of chloroplast membrane lipid metabolism in plant environmental responses. Cells 10, 706. 10.3390/cells10030706

Cutler, S.R., Rodriguez, P.L., Finkelstein, R.R., Abrams, S.R., 2010. Abscisic acid: emergence of a core signaling network. Annu. Rev. Plant Biol. 61, 651–679. 10.1146/annurev-arplant-042809-112122

de Ollas, C., Dodd, I.C., 2016. Physiological impacts of ABA–JA interactions under water-limitation. Plant Mol. Biol. 91, 641–650. 10.1007/s11103-016-0503-6

Delfosse, K., Wozny, M.R., Barton, K.A., Mathur, N., Griffiths, N., Mathur, J., 2018. Plastid envelope-localized proteins exhibit a stochastic spatiotemporal relationship to stromules. Front. Plant Sci. 9, 754. 10.3389/fpls.2018.00754

Dietz, K.-J., Turkan, I., Krieger-Liszkay, A., 2016. Redox- and reactive oxygen species-dependent signaling into and out of the photosynthesizing chloroplast. Plant Physiol. 171, 1541–1550. 10.1104/pp.16.00375

Dubiella, U., Seybold, H., Durian, G., Komander, E., Lassig, R., Witte, C.-P., Schulze, W.X., Romeis, T., 2013. Calcium-dependent protein kinase/NADPH oxidase activation circuit is required for rapid defense signal propagation. Proc. Natl. Acad. Sci. 110, 8744–8749. 10.1073/pnas.1221294110

Flors, C., Fryer, M.J., Waring, J., Reeder, B., Bechtold, U., Mullineaux, P.M., Nonell, S., Wilson, M.T., Baker, N.R., 2006. Imaging the production of singlet oxygen in vivo using a new fluorescent sensor, singlet oxygen sensor green. J. Exp. Bot. 57, 1725–1734. 10.1093/jxb/erj181

Geldner, N., Dénervaud-Tendon, V., Hyman, D.L., Mayer, U., Stierhof, Y.-D., Chory, J., 2009. Rapid, combinatorial analysis of membrane compartments in intact plants with a multicolor marker set. Plant J. 59, 169–178. 10.1111/j.1365-313X.2009.03851.x

Hare, P.D., Cress, W.A., Van Staden, J., 1998. Dissecting the roles of osmolyte accumulation during stress. Plant Cell Environ. 21, 535–553. 10.1046/j.1365-3040.1998.00309.x

Hideg, E., Kálai, T., Hideg, K., Vass, I., 1998. Photoinhibition of photosynthesis in vivo results in singlet oxygen production detection via nitroxide-induced fluorescence quenching in broad bean leaves. Biochemistry 37, 11405–11411. 10.1021/bi972890+

Ishida, T., Nagaoka, M., Akita, T., Haruta, M., 2008. Deposition of gold clusters on porous coordination polymers by solid grinding and their catalytic activity in aerobic oxidation of alcohols. Chem. Eur. J. 14, 8456–8460. 10.1002/chem.200800980

Izquierdo, Y., Muñiz, L., Vicente, J., Kulasekaran, S., Aguilera, V., López Sánchez, A., Martínez-Ayala, A., López, B., Cascón, T., Castresana, C., 2021. Oxylipins from different Pathways trigger mitochondrial stress signaling through respiratory complex III. Front. Plant Sci. 12.

Jimenez-Aleman, G.H., Jander, G., 2023. Maize defense against insect herbivory: a novel role for 9-LOX-derived oxylipins. Mol. Plant 16, 1484–1486. 10.1016/j.molp.2023.08.014

Koh, E., Chaturvedi, A.K., Javitt, G., Brandis, A., Fluhr, R., 2023. Multiple paths of plant host toxicity are associated with the fungal toxin cercosporin. Plant Cell Environ. 46, 2542–2557. 10.1111/pce.14613

Koh, E., Cohen, D., Brandis, A., Fluhr, R., 2021. Attenuation of cytosolic translation by RNA oxidation is involved in singlet oxygen-mediated transcriptomic responses. Plant Cell Environ. 44, 3597–3615. 10.1111/pce.14162

Koh, E., Fluhr, R., 2016. Singlet oxygen detection in biological systems: uses and limitations. Plant Signal. Behav. 11, e1192742. 10.1080/15592324.2016.1192742

Kwon, K.-C., Verma, D., Jin, S., Singh, N.D., Daniell, H., 2013. Release of proteins from intact chloroplasts induced by reactive oxygen species during biotic and abiotic stress. PLOS ONE 8, e67106. 10.1371/journal.pone.0067106

Li, M., Kim, C., 2023. CHAPTER One - Singlet oxygen in plants: from genesis to signaling, in: Mittler, R., Breusegem, F.V. (Eds.), Advances in botanical research, oxidative stress response in plants. Academic Press, pp. 1–42. 10.1016/bs.abr.2022.08.023

Li, X., Bao, H., Wang, Z., Wang, M., Fan, B., Zhu, C., Chen, Z., 2018. Biogenesis and function of multivesicular bodies in plant immunity. Front. Plant Sci. 9.

Marcos, R., Izquierdo, Y., Vellosillo, T., Kulasekaran, S., Cascón, T., Hamberg, M., Castresana, C., 2015. 9-Lipoxygenase-derived oxylipins activate brassinosteroid signaling to promote cell wall-based defense and limit pathogen infection. Plant Physiol. 169, 2324–2334. 10.1104/pp.15.00992

Martinière, A., Fiche, J.B., Smokvarska, M., Mari, S., Alcon, C., Dumont, X., Hematy, K., Jaillais, Y., Nollmann, M., Maurel, C., 2019. Osmotic stress activates two reactive oxygen species pathways with distinct effects on protein nanodomains and diffusion. Plant Physiol. 179, 1581–1593. 10.1104/pp.18.01065

Mattila, H., Mishra, S., Tyystjärvi, T., Tyystjärvi, E., 2023. Singlet oxygen production by photosystem II is caused by misses of the oxygen evolving complex. New Phytol. 237, 113–125. 10.1111/nph.18514

Maurel, C., Boursiac, Y., Luu, D.-T., Santoni, V., Shahzad, Z., Verdoucq, L., 2015. Aquaporins in plants. Physiol. Rev. 95, 1321–1358. 10.1152/physrev.00008.2015

Maynard, D., Chibani, K., Schmidtpott, S., Seidel, T., Spross, J., Viehhauser, A., Dietz, K.-J., 2021. Biochemical characterization of 13-lipoxygenases of *Arabidopsis thaliana*. Int. J. Mol. Sci. 22, 10237. 10.3390/ijms221910237

Miller, G., Suzuki, N., Ciftci-Yilmaz, S., Mittler, R., 2010. Reactive oxygen species homeostasis and signalling during drought and salinity stresses. Plant Cell Environ. 33, 453–467. 10.1111/j.1365-3040.2009.02041.x

Mor, A., Koh, E., Weiner, L., Rosenwasser, S., Sibony-Benyamini, H., Fluhr, R., 2014. Singlet oxygen signatures are detected independent of light or chloroplasts in response to multiple stresses. Plant Physiol. 165, 249–261. 10.1104/pp.114.236380

Murotomi, K., Umeno, A., Shichiri, M., Tanito, M., Yoshida, Y., 2023. Significance of singlet oxygen molecule in pathologies. Int. J. Mol. Sci. 24, 2739. 10.3390/ijms24032739

Nalam, V.J., Keeretaweep, J., Sarowar, S., Shah, J., 2012. Root-derived oxylipins promote green peach aphid performance on Arabidopsis foliage. Plant Cell 24, 1643–1653. 10.1105/tpc.111.094110

Nelson, B.K., Cai, X., Nebenführ, A., 2007. A multicolored set of in vivo organelle markers for co-localization studies in Arabidopsis and other plants. Plant J. 51, 1126–1136. 10.1111/j.1365-313X.2007.03212.x

Peters-Golden, M., 1998. Cell biology of the 5-lipoxygenase pathway. Am. J. Respir. Crit. Care Med. 157, S227–S232. 10.1164/ajrccm.157.6.mar4

Prasad, A., Sedlářová, M., Kale, R.S., Pospíšil, P., 2017. Lipoxygenase in singlet oxygen generation as a response to wounding: in vivo imaging in *Arabidopsis thaliana*. Sci. Rep. 7, 9831. 10.1038/s41598-017-09758-1

Pyke, K.A., 2013. Divide and shape: an endosymbiont in action. Planta 237, 381–387. 10.1007/s00425-012-1739-2

Russell, G.A., 1957. Deuterium-isotope effects in the autoxidation of aralkyl hydrocarbons. mechanism of the interaction of peroxy radicals. J. Am. Chem. Soc. 79, 3871–3877. 10.1021/ja01571a068

Sarrion-Perdigones, A., Vazquez-Vilar, M., Palací, J., Castelijns, B., Forment, J., Ziarsolo, P., Blanca, J., Granell, A., Orzaez, D., 2013. GoldenBraid 2.0: a comprehensive DNA assembly framework for plant synthetic biology. Plant Physiol. 162, 1618–1631. 10.1104/pp.113.217661

Savchenko, T., Kolla, V.A., Wang, C.-Q., Nasafi, Z., Hicks, D.R., Phadungchob, B., Chehab, W.E., Brandizzi, F., Froehlich, J., Dehesh, K., 2014. Functional convergence of oxylipin and abscisic acid pathways controls stomatal closure in response to drought. Plant Physiol. 164, 1151–1160. 10.1104/pp.113.234310

Springer, A., Kang, C., Rustgi, S., von Wettstein, D., Reinbothe, C., Pollmann, S., Reinbothe, S., 2016. Programmed chloroplast destruction during leaf senescence involves 13-lipoxygenase (13-LOX). Proc. Natl. Acad. Sci. 113, 3383–3388. 10.1073/pnas.1525747113

Sugita, C., Kato, Y., Yoshioka, Y., Tsurumi, N., Iida, Y., Machida, Y., Sugita, M., 2012. CRUMPLED LEAF (CRL) homologs of physcomitrella patens are Involved in the complete separation of dividing plastids. Plant Cell Physiol. 53, 1124– 1133. 10.1093/pcp/pcs058

Tang, G., Ma, J., Hause, B., Nick, P., Riemann, M., 2020. Jasmonate is required for the response to osmotic stress in rice. Environ. Exp. Bot. 175, 104047. 10.1016/j.envexpbot.2020.104047

Tano, D.W., Kozlowska, M.A., Easter, R.A., Woodson, J.D., 2023. Multiple pathways mediate chloroplast singlet oxygen stress signaling. Plant Mol. Biol. 111, 167–187. 10.1007/s11103-022-01319-z

Tolley, J.P., Nagashima, Y., Gorman, Z., Kolomiets, M.V., Koiwa, H., 2018. Isoform-specific subcellular localization of *Zea mays* lipoxygenases and oxo-phytodienoate reductase 2. Plant Gene 13, 36–41. 10.1016/j.plgene.2017.12.002

Vellosillo, T., Martínez, M., López, M.A., Vicente, J., Cascón, T., Dolan, L., Hamberg, M., Castresana, C., 2007. Oxylipins produced by the 9-lipoxygenase pathway in Arabidopsis regulate lateral root development and defense responses through a specific signaling cascade. Plant Cell 19, 831–846. 10.1105/tpc.106.046052

Viswanath, K.K., Varakumar, P., Pamuru, R.R., Basha, S.J., Mehta, S., Rao, A.D., 2020. Plant lipoxygenases and their role in plant physiology. J. Plant Biol. 63, 83–95. 10.1007/s12374-020-09241-x

Wang, F., Fang, J., Guan, K., Luo, S., Dogra, V., Li, B., Ma, D., Zhao, X., Lee, K.P., Sun, P., Xin, J., Liu, T., Xing, W., Kim, C., 2020. The Arabidopsis CRUMPLED LEAF protein, a homolog of the cyanobacterial bilin lyase, retains the bilin-binding pocket for a yet unknown function. Plant J. 104, 964– 978. 10.1111/tpj.14974

Wang, S., Blumwald, E., 2014. Stress-induced chloroplast degradation in Arabidopsis is regulated via a process independent of autophagy and senescence-associated vacuoles. Plant Cell 26, 4875–4888. 10.1105/tpc.114.133116

Wilson, M.E., Basu, M.R., Bhaskara, G.B., Verslues, P.E., Haswell, E.S., 2014. Plastid osmotic stress activates cellular stress responses in Arabidopsis. Plant Physiol. 165, 119–128. 10.1104/pp.114.236620

Xing, Q., Zhang, X., Li, Y., Shao, Q., Cao, S., Wang, F., Qi, H., 2019. The lipoxygenase CmLOX13 from oriental melon enhanced severe drought tolerance via regulating ABA accumulation and stomatal closure in Arabidopsis. Environ. Exp. Bot. 167, 103815. 10.1016/j.envexpbot.2019.103815

Zhang, Z.-W., Fu, Y.-F., Yang, X.-Y., Yuan, M., Zheng, X.-J., Luo, X.-F., Zhang, M.-Y., Xie, L.-B., Shu, K., Reinbothe, S., Reinbothe, C., Wu, F., Feng, L.-Y., Du, J.-B., Wang, C.-Q., Gao, X.-S., Chen, Y.-E., Zhang, Y.-Y., Li, Y., Tao, Q., Lan, T., Tang, X.-Y., Zeng, J., Chen, G.-D., Yuan, S., 2023. Singlet oxygen induces cell wall thickening and stomatal density reducing by transcriptome reprogramming. J. Biol. Chem. 299. 10.1016/j.jbc.2023.105481

